# Steroidogenic factor-1 lineage origin of skin lesions in Carney complex syndrome

**DOI:** 10.1101/2022.01.26.477839

**Authors:** Isabelle Sahut-Barnola, A-Marie Lefrancois-Martinez, Damien Dufour, Jean-Marie Botto, Crystal Kamilaris, Fabio R. Faucz, Constantine A. Stratakis, Pierre Val, Antoine Martinez

## Abstract

Carney complex (CNC) is a rare familial multi-neoplastic syndrome predisposing to endocrine and non-endocrine tumors due to inactivating mutations of *PRKAR1A* leading to perturbations of the cAMP protein kinase A (PKA) signaling pathway. Skin lesions are the most common manifestation of CNC, including lentigines, blue nevi and cutaneous myxomas, in unusual locations such as oral and genital mucosa. Unlike endocrine disorders, the pathogenesis of skin lesions remains unexplained. Here, we show that embryonic invalidation of the *Prkar1a* gene in Steroidogenic Factor-1-expressing cells, leads to the development of familial skin pigmentation alterations reminiscent of those in patients. Immunohistological and molecular analyses coupled with genetic monitoring of recombinant cell lineages in mouse skin, suggest that familial lentiginosis and myxomas occurs in skin areas specifically enriched in dermal melanocytes. In lentigines and blue nevi-prone areas from mutant mice and patients, *Prkar1a/PRKAR1A* invalidation occurs in a subset of dermal fibroblasts capable of inducing, under the influence of PKA signaling, the production of pro-melanogenic EDN3 and HGF signals. Our model strongly suggests that the origin of the typical CNC cutaneous lesions is the result of non-cell-autonomous pro-melanogenic activity of a dermal fibroblast population sharing a community of origin with SF-1 lineage.

## INTRODUCTION

Carney complex (CNC) is a rare multiple endocrine neoplasia and lentiginosis syndrome, that is familial in 70% of cases, with autosomal dominant inheritance. CNC is characterized by pigmented lesions of the skin and mucosal, myxomas (of the heart, skin and breast) and various endocrine and non-endocrine tumors. Endocrine manifestations of CNC affect the adrenal cortex (25-68% of cases), gonads (testes 35-41%, ovaries 14%), pituitary and thyroid (up to 70%). CNC patients may also present with cardiac myxoma (20-40%) and skin myxoma (20-50%). The most common clinical manifestation is abnormal skin pigmentation (60-80%) ranging from lentigines or so called freckles to blue nevi (Correa et al. 2015; Stratakis 2016; Espiard et al. 2020). Lentigines observed in CNC patients are found in unusual places such as the cheeks, vermilion border, ears, eyelids and/or genitals. In contrast to endocrine manifestations, the pathogenesis of cutaneous disorders remains unexplained. CNC is primarily caused by inactivating mutations of the *PRKAR1A* gene encoding the R1α subunit of protein kinase A (PKA), resulting in constitutive activation of cyclic adenosine monophosphate (cAMP)-PKA signaling pathway (Carney et al. 1985; Casey et al. 2000; Kirschner et al. 2000).

We have previously developed genetic mouse models carrying conditional *Prkar1a* invalidation using different Cre drivers (*Akr1b7-Cre, AS*^*Cre*^ and *Sf1-Cre*) allowing spatio-temporal R1α loss in steroidogenic cells (Sahut-Barnola et al. 2010; Drelon et al. 2016; Dumontet et al. 2018; Amaya et al. 2021). These models recapitulated the primary bilateral adrenocortical hyperplasia observed in CNC patients, associated with chronic glucocorticoid excess (Cushing’s syndrome), which resulted in metabolic comorbidities and alterations of brain structure. Remarkably, spotty skin pigmentation, the most common and earliest clinical manifestation of CNC, was only observed when *Prkar1a* was invalidated using the *Sf1-Cre* driver (*Prkar1a* ^*fl/fl*^*::Sf1-Cre*^*/+*^, named AdKO_2.0_) that was active in steroidogenic cell lineage from E10.5-E11.5 onwards (Bingham et al. 2006). Here we show that AdKO_2.0_ mice exhibit skin alterations characterized by hyperpigmentation in skin areas mirroring those of CNC patients, including lentigines, blue nevi and development of cutaneous myxomas. We used immunohistological and molecular approaches as well as cell lineage tracing studies to characterize the origin and pathogenesis of these cutaneous lesions in AdKO_2.0_ mice and compared them to skin biopsies from CNC patients. Here, we show that AdKO_2.0_ mice recapitulate skin lesions found in CNC patients and allow to identify non-cell-autonomous mechanisms involving specific dermal cell population.

## RESULTS AND DISCUSSION

### Pigmented skin lesions in AdKO_2.0_ mice are correlated with *Sf1-Cre* driver activity

The skin pigmentation phenotype of AdKO_2.0_ mice was detected at birth as a pigmented ring around the eyes. In five-day-old AdKO_2.0_ mice, hyperpigmentation was found in the perigenital skin, tail and around the nipples in female. Hyperpigmentation also formed discrete symmetrical patches under the jaw and on the joints of the upper limbs (Figure 1a-b, Figure S1a). These skin manifestations, reminiscent of lentigines, were fully penetrant and always found in these specific areas. Since adult AdKO_2.0_ mice had glucocorticoid-induced alopecia, this pattern of lentiginosis remained visible despite hair growth, as large areas around the nipples, spotty pigmentation on the scrotum and at the upper limb junctions (Figure S1b). Fontana Masson (FM) staining was carried out on skin sections from differentially pigmented areas in 5 days mice. As expected, there were no melanin pigments or histological alterations in the abdominal skin regardless of wild-type (WT) and AdKO_2.0_ mice. By contrast, melanin staining in the dermis (at least papillary and upper reticular dermis) was strongly increased in genitalia skin sections from AdKO_2.0_ mice compared to WT (Figure 1c, top panels). The specific hyperpigmentation of AdKO_2.0_ mutants does not rely on the systemic action of excess glucocorticoids, as the other two adrenal-specific *Prkar1a* invalidation models using *Akr1b7-Cre* (AdKO) or *AS*^*Cre*^ (DAdKO) drivers do not show any phenotype in the skin (Sahut-Barnola et al. 2010; Dumontet et al. 2018). This is in agreement with the observation that lentiginosis usually appears very early (at birth or during childhood) in CNC patients independently from endocrine manifestations (Stratakis 2016). Therefore, the skin phenotype of AdKO_2.0_ mice should solely be the consequence of the recombinase activity of the *Sf1-Cre* driver. To evaluate this hypothesis, we traced *Sf1-Cre* lineage in the skin by introducing the Cre recombinase *mTmG* reporter transgene (Muzumdar et al. 2007) into WT and mutant mice. In compound transgenic mice, Cre-mediated recombination fixes definitive eGFP expression in cells expressing the *Sf1-Cre* transgene and in all their descending lineages. In abdominal skin sections, eGPF immunostaining was limited to hair follicles (details in Figure S1d), while strong eGFP expression was shown in the perigenital dermis of WT and AdKO mice (Figure 1c, bottom panels). Direct eGFP fluorescence was detected in specific areas of the skin of late embryos, around the eyes, on the genitals, tail and the joint of the upper limbs (Figure S1c). This demonstrates that the skin areas where postnatal pigmentation occurs in the case of *Prkar1a* mutation, have been programmed during embryonic development and consist of specific cell lineages currently expressing or having expressed the *Sf1-Cre* transgene early. This enrichment of eGFP expression in pigmentation-prone skin areas was confirmed at the mRNA level by RT-qPCR analyses in 5-day-old WT and mutant mice (Figure 1d). Although steroidogenic activity of skin has been well documented (Slominski et al. 2013; Ceruti et al. 2018), SF-1 expression has been reported in only one study (Patel et al. 2001). Because we never detected parallel induction nor significant expression of *Sf-1* in skin samples either using RT-qPCR or immunostaining (not shown), skin eGFP-positive cells derived from *Sf-1* lineage that no longer express *Sf-1*. As expected, we showed that in AdKO_2.0_ perigenital skin, *Prkar1a* mRNA levels were decreased while transcripts for genes involved in melanosome biogenesis (*Mlana/Mart1*) and melanogenesis (*Tryp-1* and *Tryp-2*) were increased compared to WT (Figure S1e). Surprisingly, no change in mRNA levels was observed for genes encoding melanogenic transcription factors (*Mitf, Sox9, Sox10*) or dermal fibroblastic markers (*Vimentin* and *Col1a1*) (Figure S1e).

**Figure 1:**
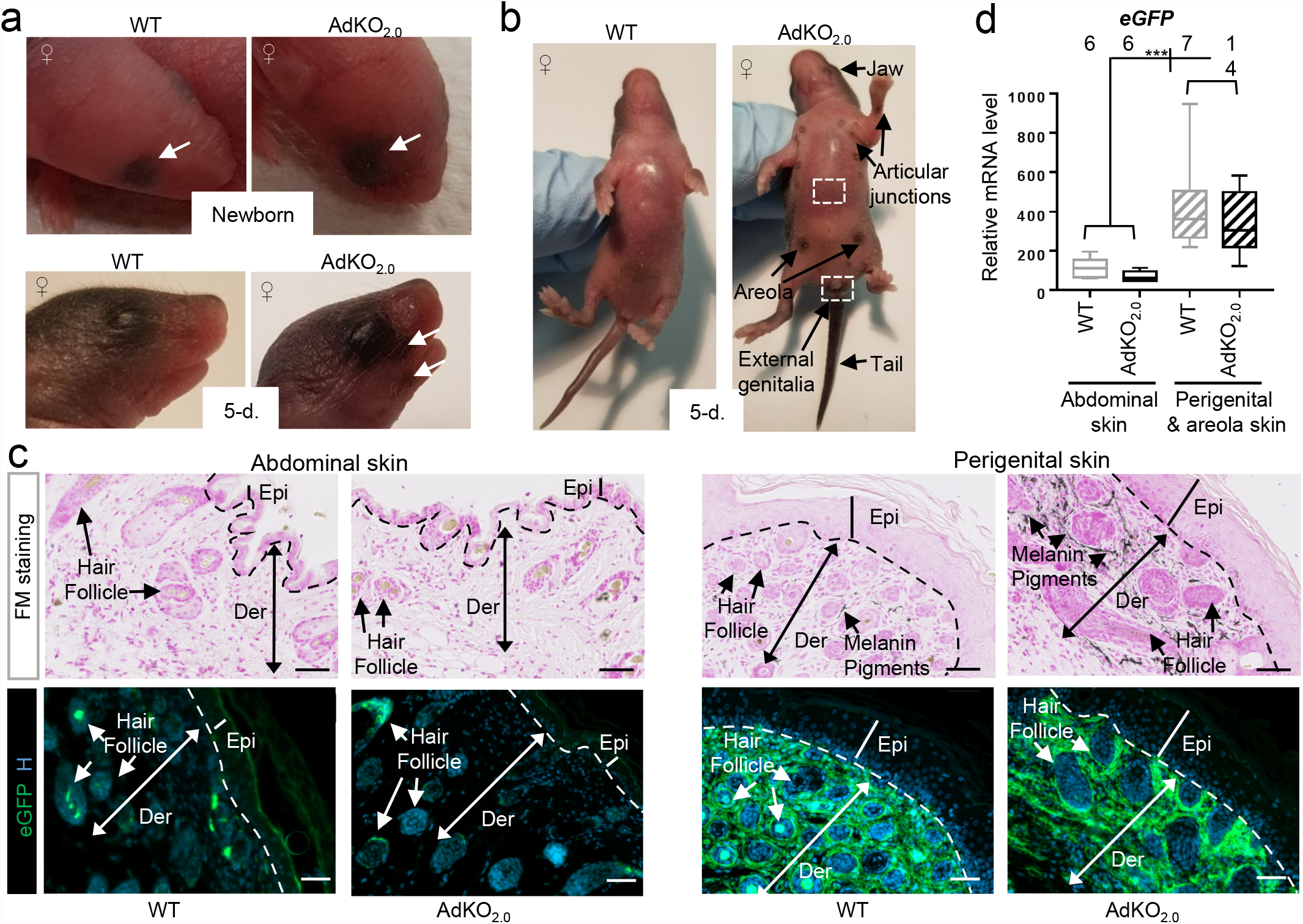
Skin pigmentation abnormalities in AdKO_2.0_ mice is associated with *Sf1-Cre* dermal activity. a-b. AdKO_2.0_ newborns (a, top) show pigmentation ring around the eye (arrows). At 5 days, pigmentation defects appear as black spots under the eye (arrows in a, bottom) and in specific locations (arrows in b, details in Figure S1). c. Fontana Masson (FM) staining (melanin) and eGFP immunodetection in WT (*Prkar1a*^*fl/+*^*::Sf1-Cre/+::R26R*^*mTmG/+*^) and AdKO_2.0_ (*Prkar1a*^*fl/fl*^*::Sf1-Cre/+::R26R*^*mTmG/+*^) (5-day-old) abdominal (top square in b) and perigenital (bottom square in b) skin sections. Hoechst nuclei staining (H). Scale bars=50μm. Epi: Epidermis, Der: Dermis. d. qPCR analysis of eGFP (proxy for recombination) in perigenital and/or areola skin compared to in WT and AdKO_2.0_ abdominal skin (5-day-old). *N*, indicated above graph. Statistical analyses: Mann Whitney test. ***P<0.001.

The lack of variation in *Mitf* expression (a major melanocyte marker and regulator of genes involved in melanogenesis) despite the marked increase in melanogenesis, has previously been described in transgenic mouse models with dermal hyperpigmentation (Wolnicka-Glubisz et al. 2013). In addition, the unaltered expression of *Mitf* in AdKO_2.0_ perigenital skin samples, was consistent with the unaltered expression of its positive upstream regulators, *Sox9* and *Sox10* (Figure 1e) (Verastegui et al. 2000; Passeron et al. 2007). Altogether these results show a direct correlation between location of lentigines in AdKO_2.0_ mice and *Sf1-Cre* activity in the dermis. *Prkar1a* inactivation in these lentigines-prone areas is expected to increase PKA activity, which in turn will enhance melanogenic activity of dermal melanocytes.

### Dermal *Sf1-Cre* lineages of the lentigines-prone areas are not melanocytes

To identify the dermal cells of the *Sf1-Cre* lineage, we carried out double immunohistochemical (IHC) analyses using on the one hand, eGFP immunostaining which marks *Prkar1a*-negative recombined cells and on the other hand, MLAN-A or VIMENTIN staining for melanocyte and fibroblast immunodetection, respectively (Figure 2a-b). In the lentigines from AdKO_2.0_ mice, there was no colocalization of eGFP and MLAN-A staining although eGFP cells were in close vicinity of MLAN-A-positive cells. In contrast, eGFP-positive cells were all positive for vimentin, indicating that *Prkar1a* invalidation occurred in dermal fibroblasts but not in dermal melanocytes.

**Figure 2:**
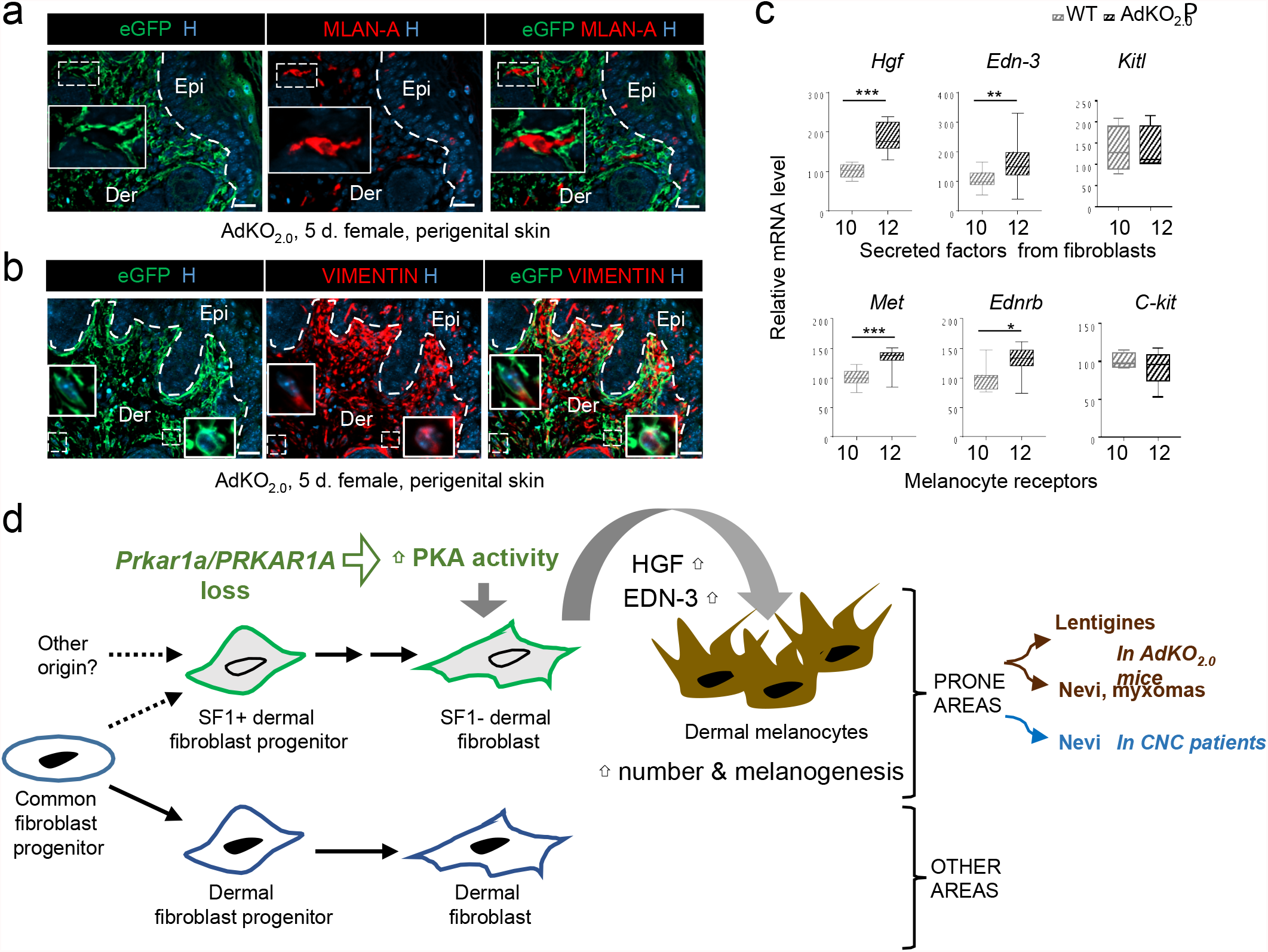
eGFP+ dermal cells (Prkar1a-KO) are PKA-reprogrammed fibroblasts producing melanogenic signals. a-b. eGFP (Prkar1a-KO cells) and MLAN-A (melanocytes) (a), or VIMENTIN (mesenchymal cells) (b) co-labelings, in AdKO_2.0_ perigenital skin (5-day-old). Hoechst nuclei staining (H). Insets, dotted frames magnification. Epi: Epidermis, Der: Dermis. Scale bars=20μm. c. qPCR analyses of WT/AdKO_2.0_ perigenital skin samples (5-day-old) for fibroblast secreted factors and corresponding receptors on melanocytes. *N*, indicated under graphs. Statistical analyses: Mann Whitney test. *P<0.05, **P<0.01, ***P<0.001. d. Schema representing skin effects of *Prkar1a/PRKAR1A* loss (AdKO_2.0_ mice/CNC patients). Increased production of diffusible factors (HGF, EDN-3) in response to PKA activation in dermal fibroblasts descending from a cell lineage that has activated *Sf1-Cre*, is expected to increase number and melanogenic activity of dermal melanocytes, favoring lentigines/nevi formation in prone areas.

Fibroblasts play an important role in regulating the activities of melanocytes by secreting a number of paracrine factors which bind to their receptors and modulate the intracellular signaling cascades linked to melanocytic functions (Wang et al. 2017). We focused on three of them and their corresponding receptors expressed on melanocytes, well characterized for their pro-melanogenic actions: hepatocyte growth factor (HGF) and MET receptor (Matsumoto et al. 1991; Kunisada et al. 2000; Kapoor et al. 2020), endothelin 3 (EDN-3) and endothelin receptor B (EDNRB) (Hosoda et al. 1994; Baynash et al. 1994; Bondurand et al. 2018) and Kit ligand (KITL) and C-KIT receptor (Geissler et al. 1988; Copeland et al. 1990; Kasamatsu et al. 2008). QPCR analyses showed an increase in mRNA levels of *Hgf* and *Edn3* and for their cognate receptors *Met* and *Ednrb*, in AdKO_2.0_ lentigines (Figure 2c). These data are consistent with previous observations from transgenic mice overexpressing HGF or EDN3 in the epidermis which displayed skin hyperpigmentation associated with an increase of dermal melanocytes (Kunisada et al. 2000; Aoki et al. 2009; Wolnicka-Glubisz et al. 2013). The mitogenic effect of HGF has also been described in primary cultures of human melanocytes (Matsumoto et al. 1991). Under physiological condition, HGF production seemed to be linked to the pigmentation of certain skin regions. Indeed, areas of the mouse skin with little hair, such as tail, nose, ear and genitalia, were shown to contain far more dermal melanocytes than hairy regions and this was associated, at least in part, with increased expression of HGF (Kunisada et al. 2000).

Our results also showed increased mRNA expression of *Edn3* and its receptor *Ebnrb*. These were in agreement with data showing that *Edn3* overexpression from E13.5, induced an increased proliferation of melanoblasts resulting in the presence of numerous dermal melanocytes and dark skin at birth (Garcia et al. 2008). Similar mechanisms involving factors identified in our animal model could also be relevant in humans. Indeed, mutations in either *EDN3* or *EDNRB* genes have been associated with diminished pigmentation in patients with Waardenburg syndrome (Mccallion and Chakravarti 2001). Furthermore, in patients with neurofibromatosis type 1, skin hyperpigmentation was associated with increased production of HGF by dermal fibroblasts (Okazaki et al. 2003).

In contrast, the mRNA encoding KITL and its receptor were not deregulated (Figure 2c). Interestingly, mouse models that overexpressed HGF or EDN3 in the skin under the control of the human *KRT14* promoter, displayed hyperpigmentation that was not influenced by KITL. Indeed, blocking KITL signaling in these mice, by using either antibodies or a genetic approach, did not affect skin hyperpigmentation while it completely inhibited that of the coat. These previous findings and our present AdKO_2.0_ model provide independent and converging arguments establishing a clear distinction between maintenance of dermal (and non-cutaneous) melanocytes and epidermal melanocytes (contributing to hair pigmentation), based on their differential requirement toward KIT signaling (Kunisada et al. 2000; Aoki et al. 2009).

In summary, the invalidation of *Prkar1a* in a subset of dermal fibroblasts (descending from a cell lineage that has activated *Sf1-Cre*) induces the expression of the pro-melanogenic factors, EDN3 and HGF, which in turn, increases the number of dermal melanocytes and/or stimulates melanogenesis in lentigines-prone areas (Model in Figure 2d). Consistent with this result, HGF is a known target of the cAMP-PKA pathway in human fibroblasts (Matsunaga et al. 1994).

### Dermal *Sf1-Cre* lineages induce nevi formation and promote the occurrence of cutaneous myxomas in AdKO_2.0_ mice

As early as 2 months of age, all AdKO_2.0_ mice developed nevi symmetrically located on the posterior aspect of the thighs. These appeared deeply embedded in the dermis and took on a blue color (Figure 3a). Histological staining and eGFP immunostaining of blue nevi sections showed that melanin pigment deposits were found in the vicinity of dermal cells that had lost *Prkar1a* conditional allele (eGFP positive) (Figure 3b). Note that clusters of sebaceous glands (positive for cytokeratin 7, not shown), formed of eGFP-negative cells with large cytoplasm, were also detected close to these pigmented formations. Double immunostaining for eGFP/MLAN-A and eGFP/VIMENTIN confirmed that *Prkar1a*-KO cells were fibroblasts (VIMENTIN positive) but not melanocytes (MLAN-A negative) (Figure S2a). As expected, qPCR analyses of nevi formed in AdKO_2.0_ mice, showed an increased expression of genes involved in melanogenesis (*Mlan-A, Tyrp-1* and *Tyrp-2*) and of fibroblastic pro-melanogenic factors (*Hgf* and *Edn3*)(Figure S2b). The association between HGF and MLAN-A protein accumulation was further confirmed in western blots (Figure 3c). This increase in local HGF should also promote the differentiation of sebaceous glands in the nevus area, which have been shown to express high levels of MET receptor (Zouboulis 2009; Wu et al. 2014). Our results indicate that as with lentigines described above, blue nevi formation in AdKO_2.0_ mice arises from activation of a pro-melanogenic fibroblast population, following *Prkar1a* loss (Figure 2d and see Figure 4a for a summary diagram of pigmented skin areas in AdKO_2.0_ mice).

**Figure 3:**
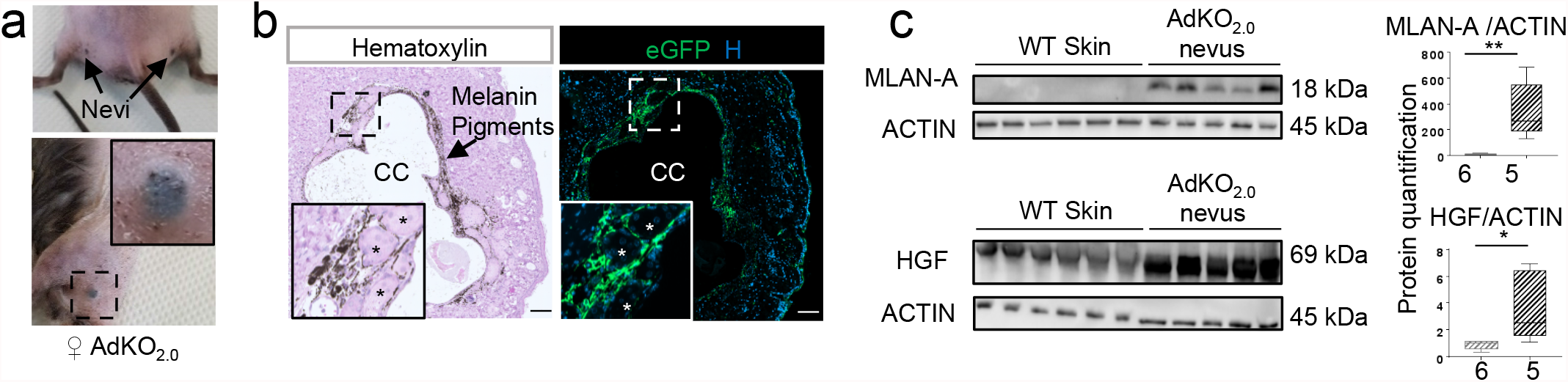
Increased PKA activity is associated with blue nevi in AdKO_2.0_ skin. a. View of symmetrical nevi (top) and magnification (bottom) in 4-6-month-old females. b. Hematoxylin staining and eGFP immunodetection of blue nevus section from 6-month-old AdKO_2.0_ female. In nevus section, the central cavity (CC) represents the place where melanin accumulates (eliminated by histological treatments). The periphery of the cavity is lined by melanin pigments, eGFP-positive cells and by clusters of sebaceous glands formed of eGFP-negative cells with large cytoplasm (* in the magnification panels). Scale bars=100μm. c. Western blot analyses of MLAN-A and HGF protein accumulation in skin extracts from WT and AdKO_2.0_ (nevi) mice. Graphs showed quantification normalized to ACTIN signal. *N*, indicated under graphs. Statistical analyses: Mann Whitney test. *P<0.05; **P<0.01.

**Figure 4:**
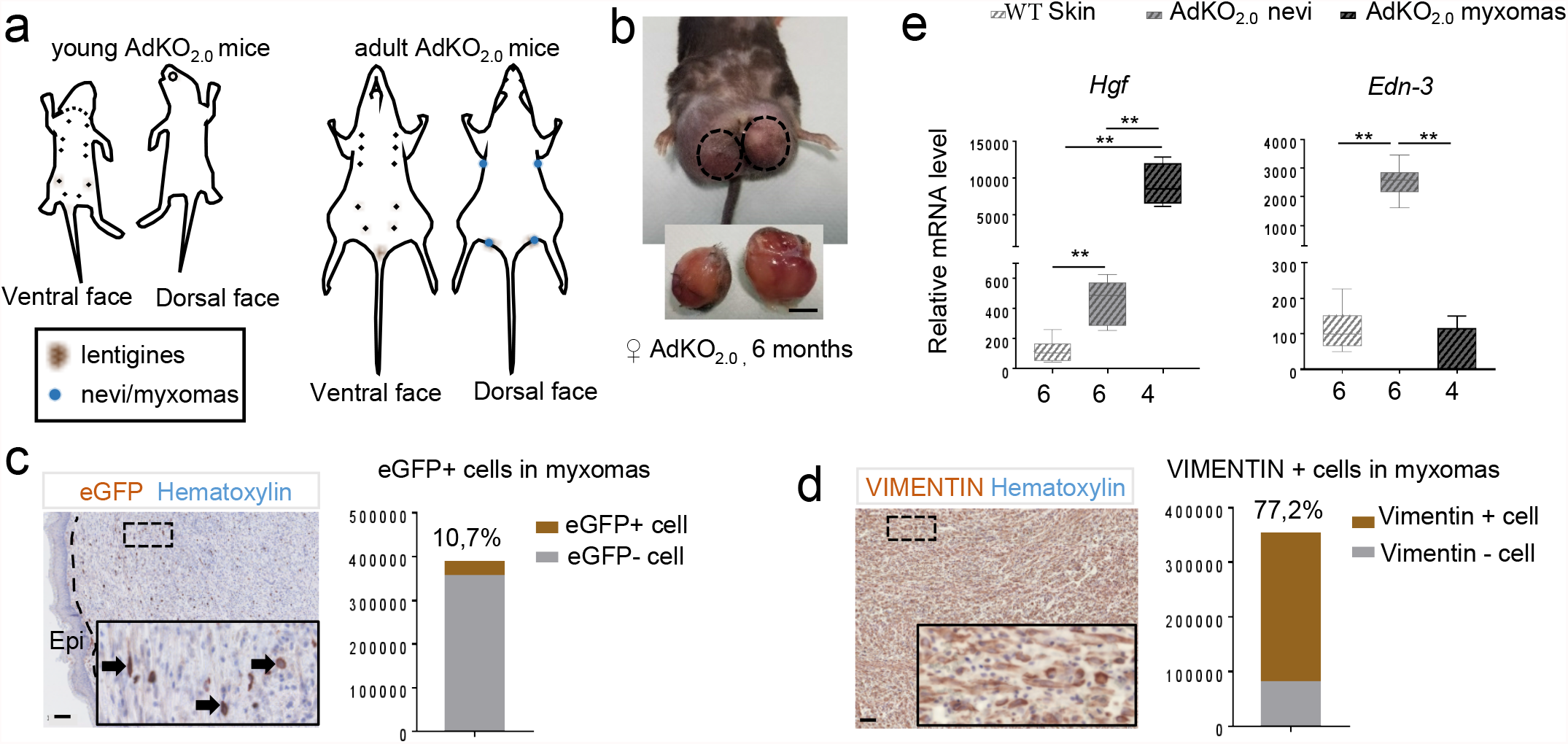
Cutaneous myxomas in AdKO_2.0_ mice. a. Skin lesions maps in 5-day- and 2-month-old mice. Head lentigines are no longer seen in adults due to hair growth. Nevi form symmetrically on back of thighs and shoulders. b. Myxomas emerge in 44% of AdKO_2.0_ mice (≥6 months), in areas where nevi form. Scale bar=1cm. c-d. The tumor is delimited by epidermis (Epi). eGFP (c) or Vimentin (d) immunodetections (insets, frame magnification) and hematoxylin counterstaining. Percentages of eGFP-positive (Prkar1a-KO) and Vimentin-positive cells are calculated from counts of four different tumors. Scale bar=100μm. e. qPCR analyses of fibroblastic secreted factors in WT skin (sampling located where nevi form in mutants) and AdKO_2.0_ nevi (3-month-old females) and in AdKO_2.0_ myxomas. *N*, indicated under graphs. Statistical analyses: Mann Whitney test. **P<0.01.

In 20-50% of CNC patients, cutaneous myxomas were found localized in specific areas such as the eyelid, external ear canal, areola and genitals (Correa et al. 2015; Espiard et al. 2020). About 44% of AdKO_2.0_ mice from 6 months of age, developed large skin tumors localized either on the thighs or on the shoulder joint (where nevi occur) (Figure 4b). The tumor masses always appeared dense, poorly vascularized, non-melanic and encapsulated. Hematoxylin staining showed an epidermal delineation of the tumor (Figure 4c). CNC skin myxomas are composed of spindle-shape cells and polygonal cells (Hachisuka et al. 2006). Similar histological features were also observed in skin tumor masses of AdKO_2.0_ mice. The eGFP-labeled cells (*Prkar1a*-KO cells) were found scattered throughout the entire tumor and accounted for at least 10% of the total cell population, which consisted mainly of mesenchymal cells (vimentin positive), indicating the myxomatous nature of these cutaneous tumors (Figure 4d). QPCR analyses suggested that formation of skin myxomas could involve the dramatic increase (x100) in *Hgf* gene expression while, as expected, *Edn-3* mRNA levels were not increased in these nonpigmented lesions compared to blue nevi (Figure 4e). In conclusion, AdKO_2.0_ mice develop cutaneous tumors that are hallmarks of skin lesions found in CNC patients. The occurrence of myxomas in sites where blue nevi initially develop, suggests a possible continuity between these pigmented lesions and myxoma formation.

### Lentigines and blue nevi from CNC patients

With access to some rare skin samples from CNC patients, we characterized the sites of melanin accumulation using Fontana Masson or hematoxylin staining (Figure 5a, top; Figure S3a, top). Lentigines (Car-lentigo-1, Car-lentigo-2) showed high levels of melanin pigments in the basal layer of the epidermis as previously described (Courcoutsakis et al. 2013), while the pigmentation in blue nevi (Car-blue nevus-3, Car-blue nevus-4) was rather concentrated in the dermis. In normal skin and lentigines, HGF immunostaining was present in the epidermis and in scattered cells in the dermis, in agreement with the literature showing HGF production in keratinocytes and fibroblasts (Serre et al. 2018).

**Figure 5:**
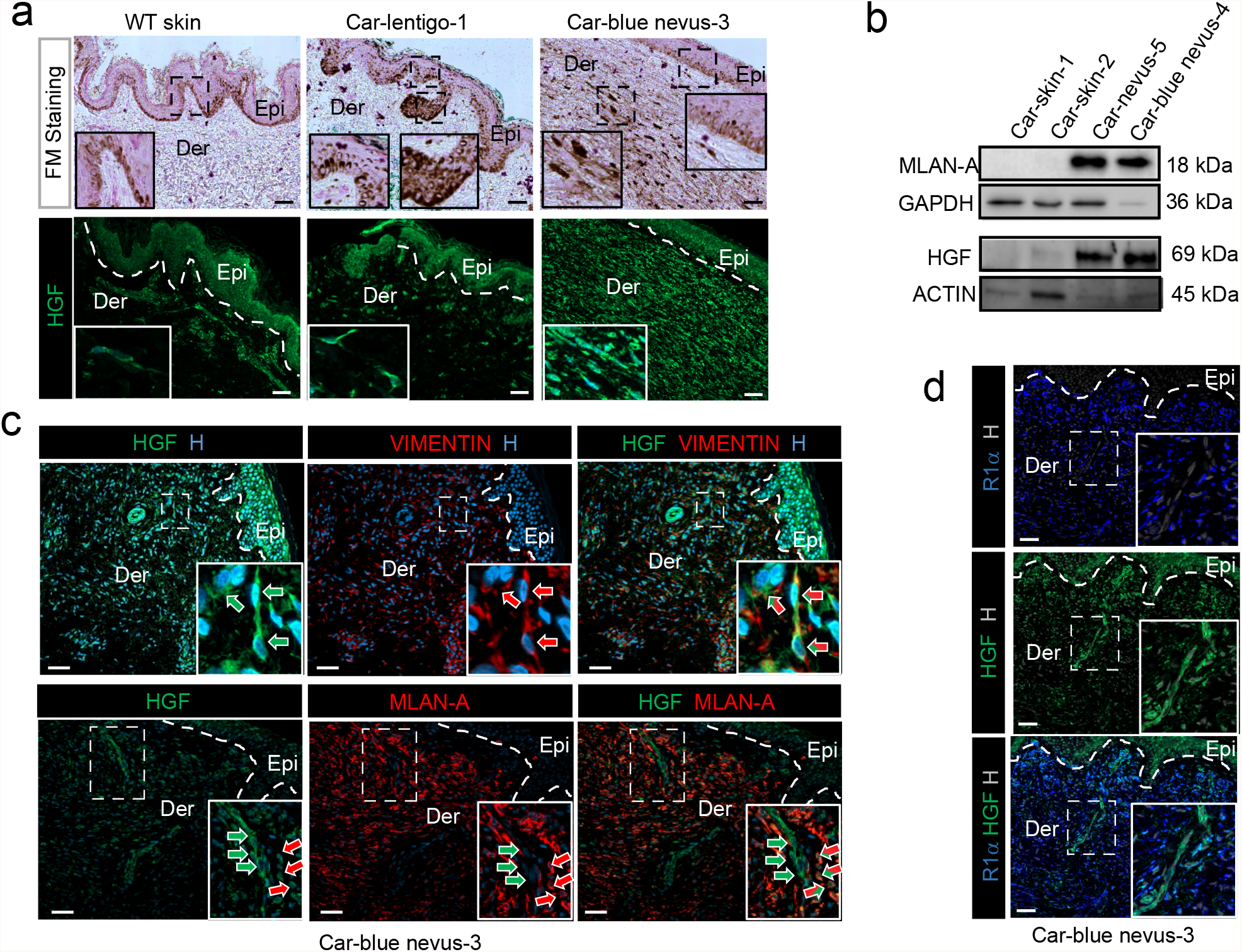
Increased HGF is associated with dermal melanin in blue nevi from CNC patients. a. Fontana Masson (FM) staining and HGF immunodetection of normal human skin and lentigines/blue nevi from CNC patients. b. Western blot analysis of MLAN-A and HGF protein accumulation in skin extracts from CNC patients. c. Co-immunolabeling of HGF (green arrows) and VIMENTIN (mesenchymal cells, red arrows) in top panels, or MLAN-A (melanocytes, red arrows) in bottom panels in blue nevus from CNC patients (Striped arrows, cells co-labelled with HGF and VIMENTIN/MLAN-A). d. Co-immunofluorescent labeling of R1α and HGF in blue nevus from CNC patients. Hoechst nuclei staining (H). Insets are magnification of dotted frames. Epi: Epidermis, Der: Dermis. Scale bars=50μm.

In contrast, in blue nevi, HGF labeling was strong in the dermis (Figure 5a, bottom; Figure S3a, bottom). Western blot analyses confirmed that HGF contents were high in pigmented skin lesions from CNC patients that expressed high levels of the melanosome biogenesis marker MLAN-A (Figure 5b).

HGF/VIMENTIN and HGF/MLAN-A double immunostaining was performed in blue nevi sections (Figure 5c, similar results for Car-blue nevus-4, not shown). These experiments showed co-localization of HGF staining with most of the numerous dermal fibroblasts (VIMENTIN-positive). Using HGF/MLAN-A co-staining, two cell types could be identified:

1) strongly HGF-positive/MLAN-A-negative cells that should be HGF producing fibroblasts, and 2) double positive cells that should be dermal melanocytes, on which HGF binds to enhance melanogenesis.

We then interrogated the expression pattern of *PRKAR1A* in CNC skin samples. As expected, in CNC adrenal gland, R1α immunostaining was only detected in internodular cortex where *PRKAR1A* is expressed from the intact allele and absent from nodules due to loss of heterozygosity (Figure S3c). In normal skin or CNC lentigines, R1α staining was detected in both the dermis and in epidermal melanocytes (MLAN-A positive cells) (Figure S3b, bottom right). In blue nevi, R1α staining was present in most dermal cells, with the exception of a few clusters of fibroblasts that strongly expressed HGF, presumably due to increased PKA activity (Figure 5d). The lentigines formation in CNC patients did not appear associated with an increase in HGF (Figure 5a, Figure S3a). This discrepancy with the AdKO_2.0_ model could rely on species specificity, because unlike humans, the adult mouse epidermis lacks melanocytes and, therefore, skin pigmentation solely relies on dermal melanocytes.

In conclusion, our results strongly argue that loss of R1α in some dermal fibroblasts induces PKA-dependent production of secreted growth factors including HGF, which in turn, stimulate melanogenic activity of dermal melanocytes. This non-cell-autonomous mechanism of *PRKAR1A*-dependent blue nevus pathogenesis in CNC patients is consistent with what we demonstrated in the skin lesions of AdKO_2.0_ mice, which ultimately might evolve into massive skin myxomas (Figure 2d). There is great heterogeneity in the origin, location and functions of skin fibroblasts. For example, cell fate mapping analyses performed on the avian or mouse embryo revealed that skin fibroblasts of the dorsal, ventral, lateral or craniofacial dermis are derived from different embryonic tissues (Thulabandu et al. 2018). Our cell lineage tracing experiments in mice suggest that the pigmented skin lesions typical of CNC, form through the *Prkar1a*-dependent activation of a specific subpopulation of dermal fibroblasts sharing common origin with the steroidogenic lineage. Most recent studies investigating the lineage diversity of fibroblast populations have focused primarily on the mechanisms of hair follicle formation and repair (Driskell et al. 2013; Joost et al. 2020). Therefore, we hypothesize that the specific distribution of cutaneous pigmented lesions in CNC may rely on the paracrine activity of specific dermal fibroblast populations, preferentially located in areas enriched in dermal melanocytes, sharing possible common origins with SF-1 lineage and having increased sensitivity to changes in PKA activity.

## MATERIALS & METHODS

### Mouse

The *Prkar1a*^*fl/fl*^*::Sf1*^*cre/+*^*::R26R*^*mTmG/+*^ mouse line have been described previously ((Muzumdar et al. 2007; Drelon et al. 2016). Mice were all maintained and bred on a mixed background. Throughout, WT refers *Prkar1a*^*fl/+*^*::Sf1*^*cre/+*^*::R26R*^*mTmG/+*^ in which we did not observe any discernible phenotype. Littermate control animals were used in all experiments.

### Histology and immunostaining

For general morphology and melanin detection, sections were stained with hematoxylin and eosin and with Fontana Masson staining (MF-100-T ; Biognost), following the manufacturer’s instructions. Immunohistological methods have been previously described (Drelon et al. 2016). Cell count was performed using the QuPath software (Bankhead et al. 2017). See Table S1 for conditions used.

### Western blot

Twenty five (mouse WB) or twelve (human WB) micrograms of total proteins were loaded on 4/15% SDS-page gel, transfer on nitrocellulose and detected with primary antibodies (Table S1). Signals were quantified with ChemiDoc MP Imaging System camera system (Bio-rad) and Image Lab software (Bio-rad). Expression of proteins were normalized to expression of the ACTIN or GAPDH protein.

### Real-time PCR analysis

Total RNAs were extracted from skin using the RNAII nucleotide extraction kit (Macherey Nagel), following the manufacturer’s instructions. One microgram of total mRNAs was reverse transcribed for 1h at 37C with 5pmol of random hexamers primers, 200 units reverse transcriptase (M-MLV RT, M1701, Promega), 2mM dNTPs and 20 units RNAsin (N2615, Promega). One microlitre of a one-fourth dilution of cDNA was used in each qPCR. PCR reactions were conducted with SYBR qPCR Premix Ex Taq II Tli RNaseH+ (TAKRR820W,Takara). Primer pairs are listed in Table S2. For each experiment and primer pairs, efficiency of PCR reactions was evaluated by amplification of serial dilutions of a mix of cDNAs. Relative gene expression was obtained by the DDCt method after normalization to actin.

### Samples from patients

Histological slides of skin lesions from CNC patients and skin biopsies were obtained from Pr C.A. Stratakis lab, NIH. Human adult normal skin sections were used as control for histological analyses (UM-HuFPT136, CliniSciences). Sample information are available in Tables S3 & S4.

### Statistics

Statistical analyses were conducted by Man Whitney test. P values of less than 0.05 were considered significant. All the data are represented by Box & Whiskers Plots with the minimum and maximum value and the median.

### Study Approval

All animal studies were approved by Ethics Committee for Animal Experimentation in Auvergne (N° 21153-2019061912044646V2) and were conducted in agreement with international standards for animal welfare.

All patients gave informed consent for the use of their resected tissues for research purposes.

## CONFLICT OF INTEREST

The authors have declared that no conflict of interest exists

## ACKNOWLEDGMENTS

This work was funded through institutional support from Centre National de la Recherche Scientifique, INSERM, Université Clermont-Auvergne and by Agence Nationale pour la Recherche (ANR-18-CE14-0012-02).

## AUTHORS CONTRIBUTIONS

ISB, AMLM, DD, JMB and AM designed experiments. ISB, AMLM, PV and AM prepared manuscript. ISB and AM supervised the project. CK, FRF and CAS provided human samples and clinical information. All authors edited the manuscript.

## FIGURE LEGENDS

**Table S1:**
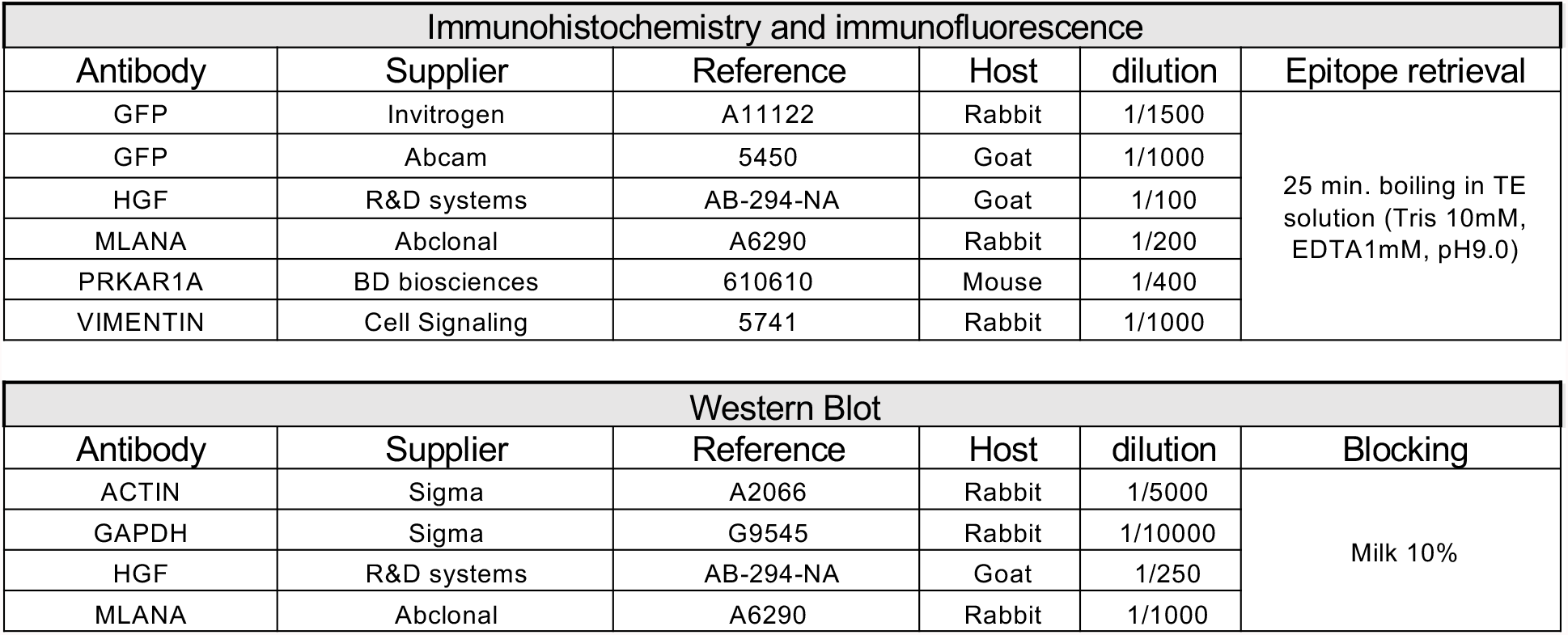
Immunohistological and western blot conditions

| Immunohistochemistry and immunofluorescence |  |  |  |  |  |
| --- | --- | --- | --- | --- | --- |
| Antibody | Supplier | Reference | Host | dilution | Epitope retrieval |
| GFP | Invitrogen | A11122 | Rabbit | 1/1500 | 25 min. boiling in TE solution (Tris 10mM, EDTA1mM, pH9.0) |
| GFP | Abcam | 5450 | Goat | 1/1000 |  |
| HGF | R&D systems | AB-294-NA | Goat | 1/100 |  |
| MLANA | Abclonal | A6290 | Rabbit | 1/200 |  |
| PRKAR1A | BD biosciences | 610610 | Mouse | 1/400 |  |
| VIMENTIN | Cell Signaling | 5741 | Rabbit | 1/1000 |  |

| Western Blot |  |  |  |  |  |
| --- | --- | --- | --- | --- | --- |
| Antibody | Supplier | Reference | Host | dilution | Blocking |
| ACTIN | Sigma | A2066 | Rabbit | 1/5000 | Milk 10% |
| GAPDH | Sigma | G9545 | Rabbit | 1/10000 |  |
| HGF | R&D systems | AB-294-NA | Goat | 1/250 |  |
| MLANA | Abclonal | A6290 | Rabbit | 1/1000 |  |

**Table S2:**
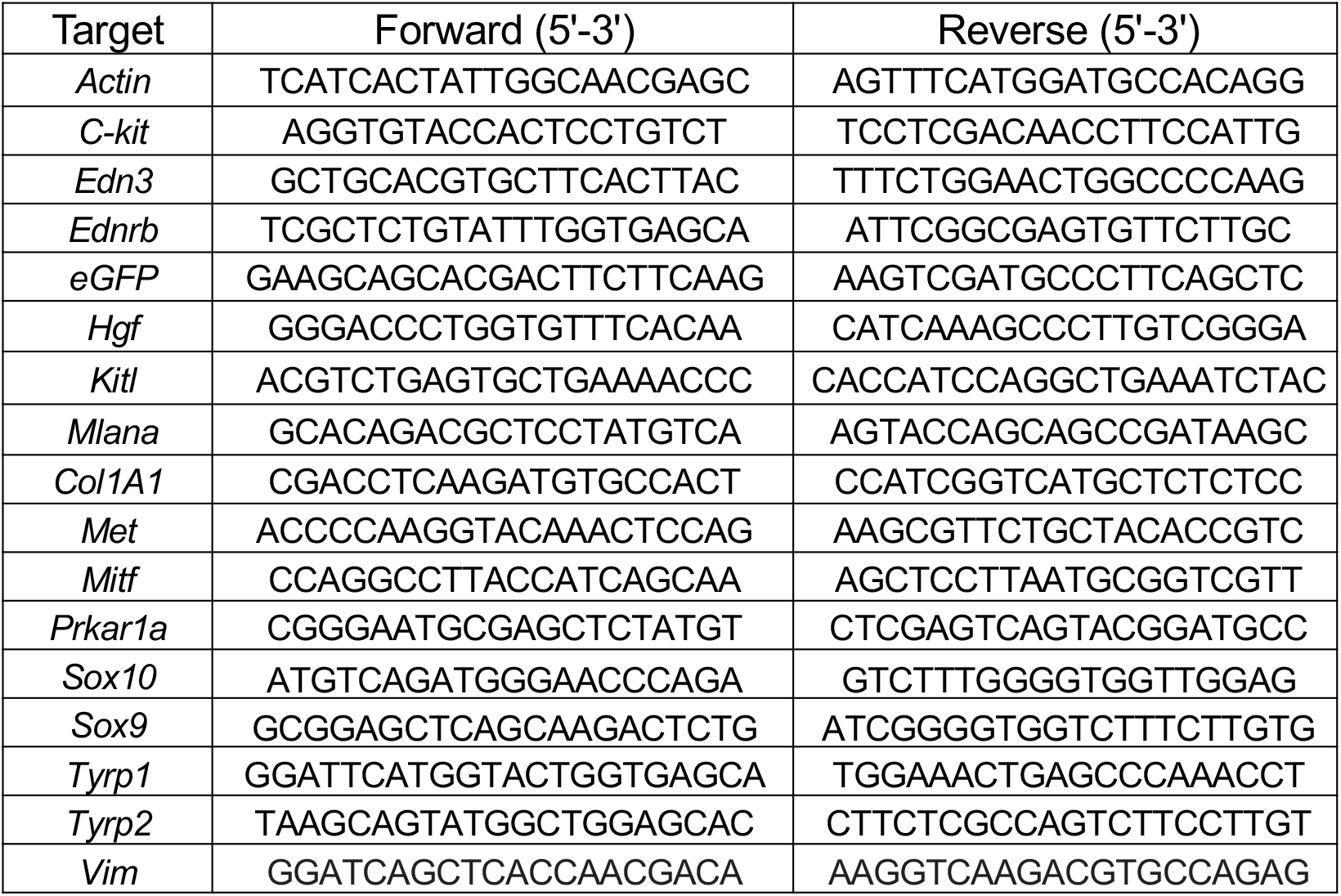
Sequences of the primers used for RTqPCR

**Table S3:** Human skin Information

| Number | Identification Number | Diagnosis | Sample |
| --- | --- | --- | --- |
| Car-skin-1 | CAR 27.04 | Skin myxoma | Sample for WB |
| Car-skin-2 | CMC.01 | Spitz nevus | Sample for WB |
| Car-nevus-5 | CAR46.01 | Nevus | Sample for WB |
| Car-nevus-4 | CAR 20.15 | Blue nevus | Sample for WB and Histological slides |
| Car-nevus-3 | CAR 742.01 | Blue nevus | Histological slides |
| Car-lentigo-1 | CAR 58.02 | Benign lentigo | Histological slides |
| Car-lentigo-2 | CAR 510.01 | Benign lentigo | Histological slides |

**Table S4:** Clinical data of CNC patients

|  | PRKAR1A mutation | Age | Sex | CNC associated manifestations | Hormonal data |
| --- | --- | --- | --- | --- | --- |
| CAR27.04 | c.124C>T/p.Arg42X | Data not available | Female | Data not available | Data not available |
| CMC.01 | Suspected somatic defect | Data not available | Male | Data not available | Data not available |
| CAR46.01 | No mutation | 46 y | Male | GH excess, thyroid nodules, LCCSCT, PPNAD, nevus | IGF-1: 410mg/mL (110-267)<br>Free T4: 0.9ng/dL (0.7-1.5)<br>TSH: 0.98uIU/mL (0.4-4)<br>Testosterone: 274ng/dL (240-871)<br>Midnight serum cortisol: 3ug/dL<br>Prolactin: 2.5 ug/L (1-25)<br>GH at 2 hours of OGTT: 2.8ng/mL<br>Urinary free cortisol: 8ug/24h,4.8ug/24h, 6.7ug/24h, and 6.8ug/24h (3.5-45) |
| CAR20.15 | c.491_492delTG/p.Val164fsX4 | 32 y | Female | lentiginos, blue nevi, growth hormone excess, PPNAD, breast myxomas, pituitary adenoma, breast ductal adenoma, ovarian cyst | Midnight cortisol: 3.3 mcg/dL<br>TSH 1.44 uIU/mL (0.4-4)<br>Free T4 0.8 ng/dL (0.7-1.5)<br>DHEAS 1.64 mcg/mL (18-244)<br>GH at 2 hours of OGTT: 4.9ng/mL<br>IGF-1 486ng/mL (110-267)<br>FSH 4 U/L (Follicular 3-11, Midcycle 6-21, Luteal 1-9)<br>LH <1 U/L Follicular 1-12 , Midcycle 17-77, Luteal 0-15) |
| CAR742.01 | No mutation |  | Male | lentiginos, blue nevus, pigmented epithelioid melanocytoma.<br>Patient also has a history of multiple myeloma. | Data not available |
| CAR58.01 | Exon 2 Deletion | 43 y | Female | benign lentigo, cardiac myxoma, PPNAD, hyperprolactinemia, thyroid nodules, lentiginos | Midnight serum cortisol 7.2mcg/dL<br>FSH 7U/L (Follicular 3-11, Midcycle 6-21, Luteal 1-9)<br>TSH 1.27mIU/mL (0.4-4)<br>DHEAS <15ug/dL (18-244)<br>Estradiol 79 pg/mL (15-350)<br>ACTH 25.5 pg/mL (9-52)<br>Thyroxine 8.4mcg/dL (4.5-12.5)<br>LH 4 U/L Follicular 1-12 , Midcycle 17-77, Luteal 0-15)<br>IGF-1 117ng/mL (110-267)<br>Prolactin 30.9 mcg/L (1-25)<br>GH at 2 hours of OGTT 0.3ng/mL |
| CAR510.01 | No mutation | 57 y | Female | benign lentigo, pituitary adenoma, GH excess, hyperprolactinemia, lentiginos. Patient also has a history of infiltrating ductal breast carcinoma and a schwannoma. | IGF-1 (post-TSS): 218ng/mL (110-267)<br>Estradiol (post-TSS): 18 pg/mL (15-350)<br>LH (post-TSS): 2 U/L (11-40)<br>Free T4 (post-TSS): 1.2ng/dL (0.7-1.5)<br>TSH (post-TSS): 0.95 uIU/mL (0.4-4)<br>GH at 2 hours of OGTT (prior to TSS): 6.3ng/mL<br>ACTH (post-TSS): 7pg/mL (9-52)<br>FSH (post-TSS): 9U/L (22-153)<br>Prolactin (post-TSS): 11ug/L (1-25)<br>Midnight serum cortisol (post-TSS): 1.4mcg/dL<br>UFC: 52mcg/24h and 25mcg/24h |

**Figure S1:**
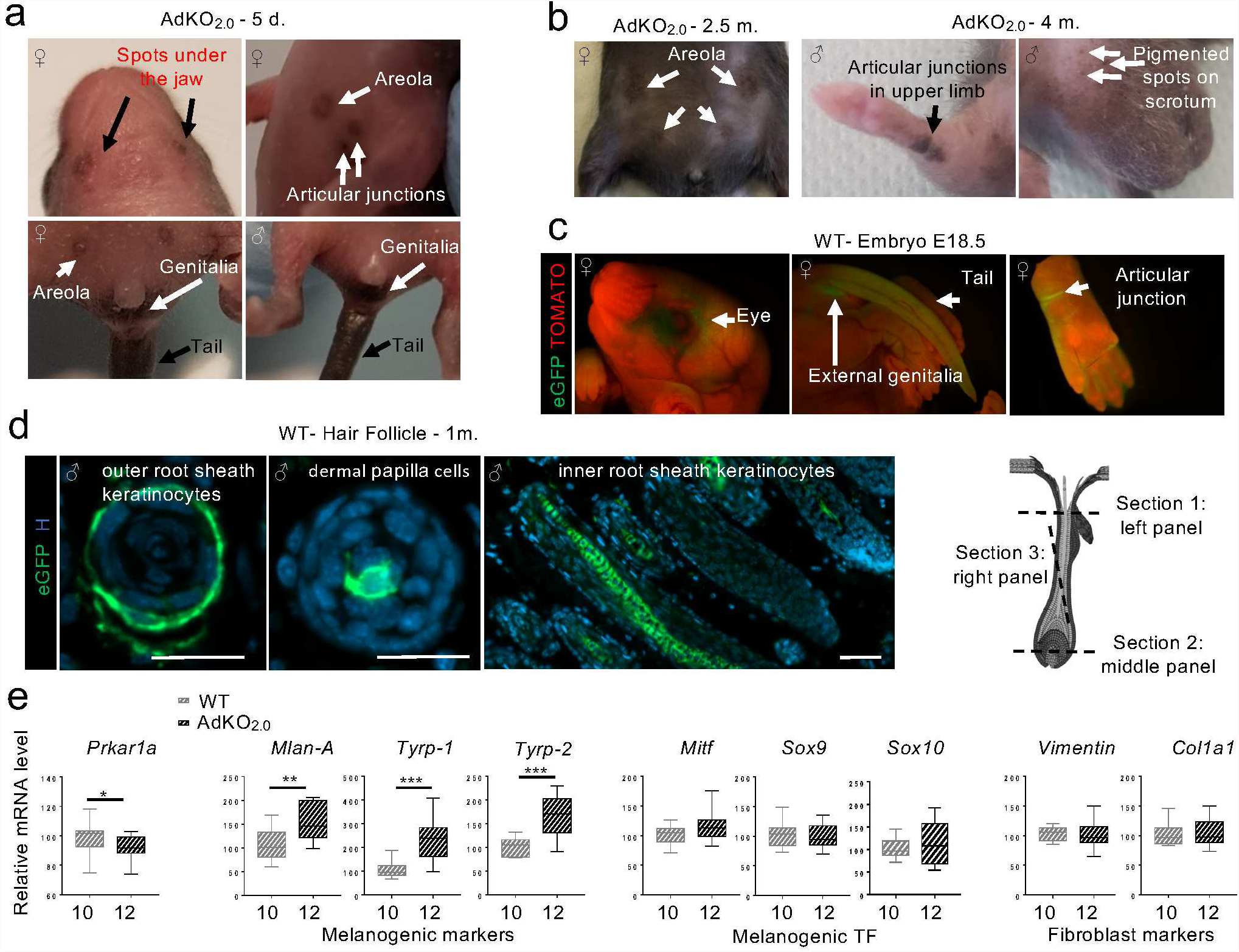
Sf1-Cre activity in AdKO_2.0_ and WT mice. a-b Pigmentation in young (a) and adult AdKO_2.0_ mice (b). c. *Prkar1a*^*fl/+*^ *::Sf1-Cre/+::R26R*^*mTmG/+*^ (referred to as WT) mice allow tracking eGFP expression sites that report *Sfl-Cre* driver activity. In E18.5 embryos, direct eGFP fluorescence highlights Cre cutaneous activity (arrows), d In various skin types, Cre is also active in hair follicles: representative picture from one-month-old WT male skin section (follicle drawing inspired, from Slominski, et al. 2013). Scale bars=50µm. e. qPCR analyses from W1 and AdKO_2.0_ perigenital skins (5-day-old) confirm melanogenic markers increased expression. As expected, regarding restricted Cre activity, *Prkarla* mRNA level is only slightly decreased in AdKO_2.0_. mRNAs expression of melanogenic transcription factors (TF) or fibroblast markers is unchanged. *N*, indicated under graphs. Statistical analyses: Mann Whitney test. *P<0.05, **P<0.01, ***P<0.001.

**Figure S2:**
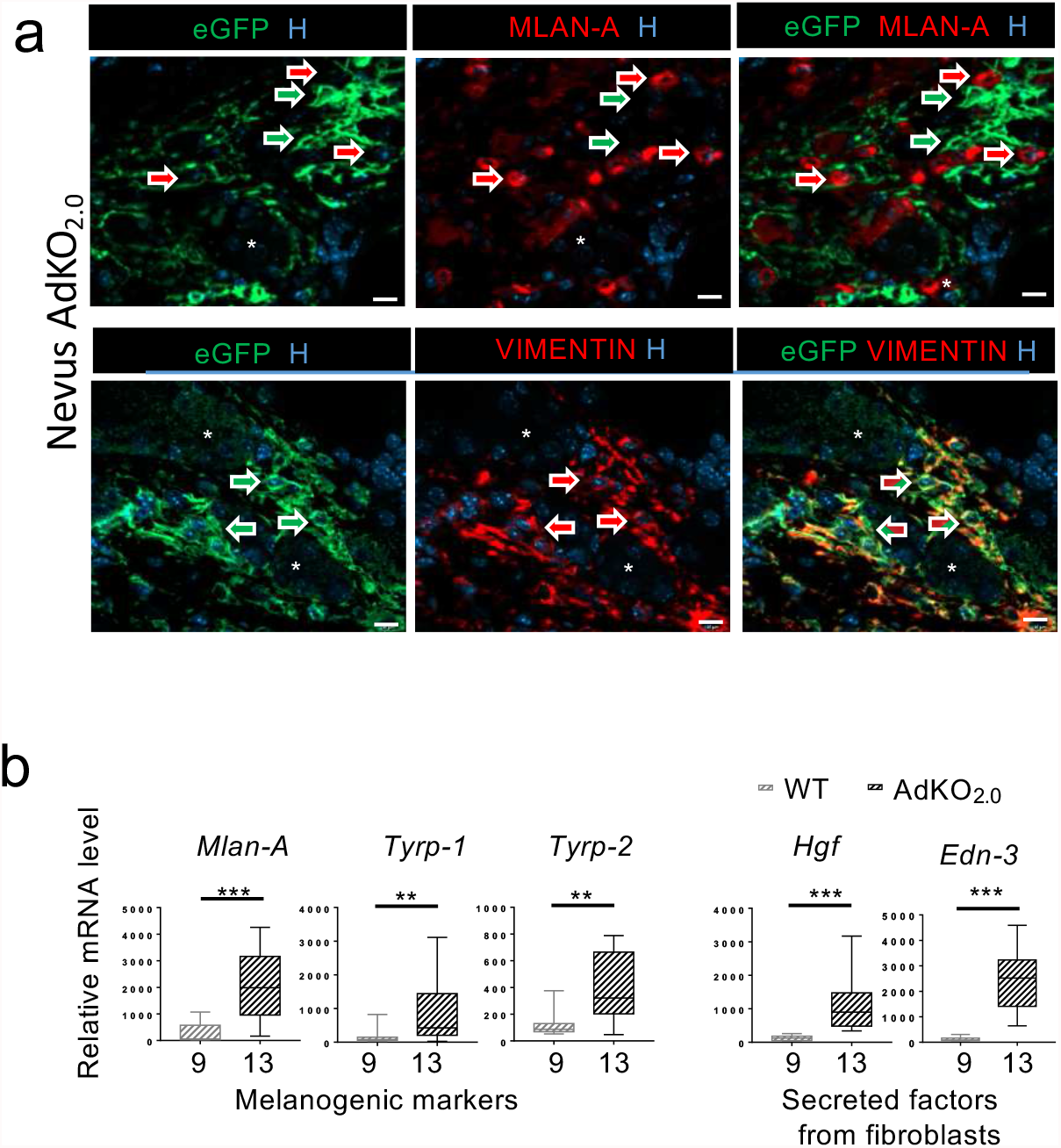
PKA increase affects dermal fibroblasts in AdKO_2.0_ blue nevi. a. Co-immunofluorescent labeling of eGFP (Prkar1a-K0 cells, green arrows) and Mlan-A (melanocytes, red arrows) in top panels, or Vimentin (mesenchymal cells, red arrows) in bottom panels, in the blue nevus shown in Figure 3a. Cells co-labelled with eGFP and Vimentin are identified by striped arrows. Hoechst nuclei staining (H). Sebaceous glands (*). Scale bars=10µm. b. qPCR analyses of melanogenic markers and fibroblastic secreted factors in WT skin (sampling located where nevi form in mutants) and AdKO_2.0_ nevi from 3-4-month-old females. *N*, indicated under graphs. Statistical analyses: Mann Whitney test. **P<0.01, ***P<0.001.

**Figure S3:**
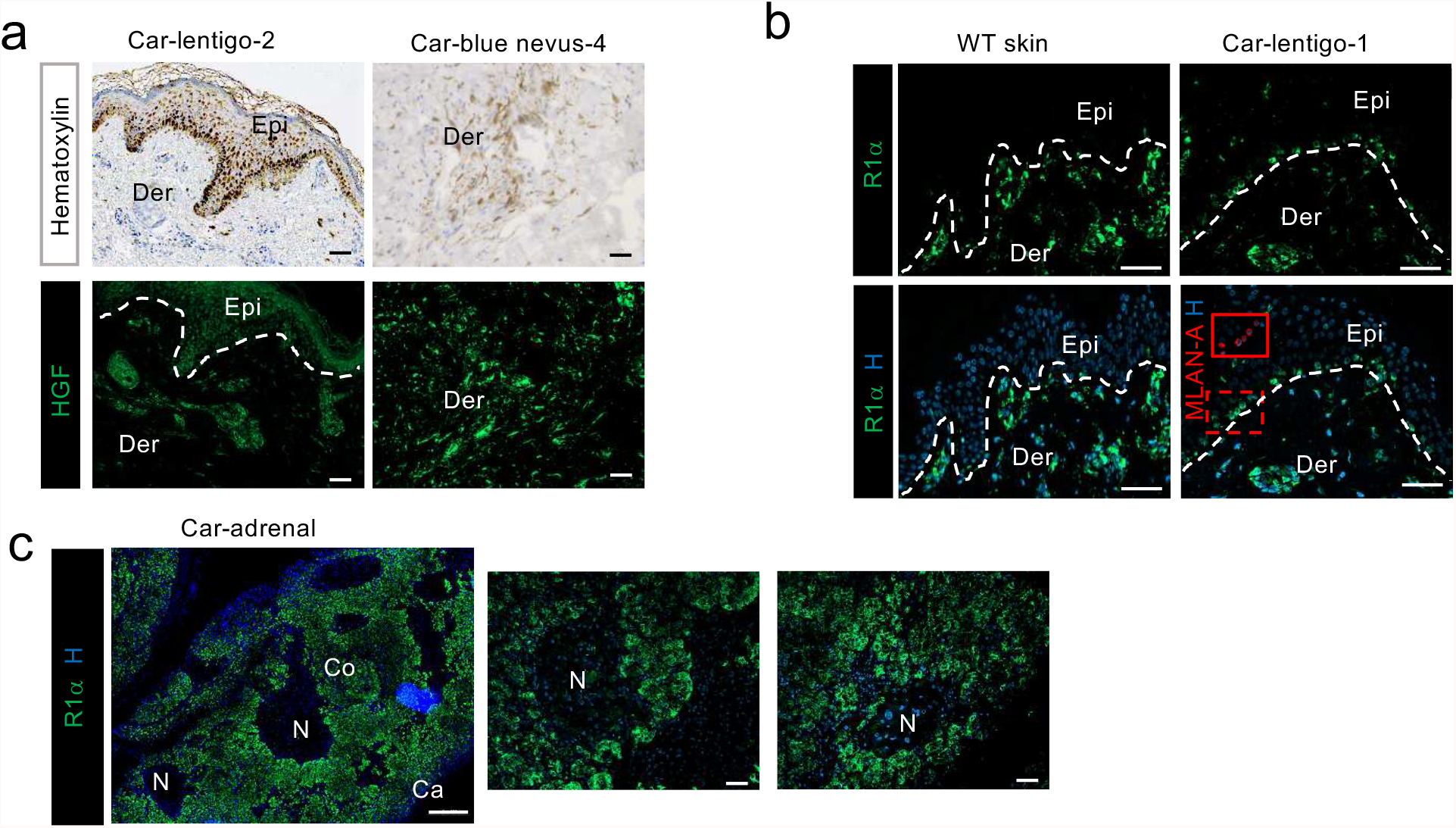
Lentigines, blue nevi and adrenal glands in CNC patients, a. Hematoxylin staining (black staining is melanin pigments) and immunostaining of HGF in lentigines and in blue nevus from patients with Carney complex. Scale bars=50µm. b. Immunofluorescent labeling of R1α in lentigines from CNC patients. Hoechst nuclei staining (H). The red inset (corresponding to dotted red frame) shows immunolabeling of MLAN-A. Epi: Epidermis, Der: Dermis. Scale bars=50µm. c. Immunolabeling of R1α in micronodular hyperplastic adrenal gland from CNC patients. Nodules (N) are negative for R1α, confirming the specificity of the antibody. Ca, capsule; Co, cortex. Scale bars=500pm in the left panel and =50pm in the 2 right-hand panels.

## Notes

### Competing Interest Statement

The authors have declared no competing interest.

